# Local RhoA activation induces septin recruitment

**DOI:** 10.1101/2025.09.29.679255

**Authors:** Shreya Chandrasekar, Margaret E. Utgaard, Bradley Somerfield, Huini Wu, Jordan R. Beach, Patrick W. Oakes

## Abstract

The regulation of the actin cytoskeleton is key for controlling cell shape and structure. While the Rho GTPase RhoA is well known to regulate the actomyosin cytoskeleton, its function in controlling the septin cytoskeleton remains unclear. As RhoA interactions can vary in both time and space, they can be challenging to discern from traditional bulk biochemical assays. Here we use multiple optogenetic tools to spatially and temporally increase myosin localization, stimulate contractile force, and activate RhoA, to investigate how RhoA and its downstream effector myosin impact the septin cytoskeleton. We find that neither local accumulation of myosin nor increased activity of myosin is sufficient to alter septin architecture. Local activation of RhoA, however, results in a local increase in septin accumulation. Importantly, this septin increase is independent of the scaffolding protein anillin, which can directly bind both septin and RhoA. Together these data expand the potential role of septins in mediating RhoA signaling by stimulating the remodeling of the septin cytoskeleton.

## Introduction

The cytoskeleton plays a key role in determining cell shape, which in turn is intimately connected to cell behavior. Cytoskeletal networks, in conjunction with their associated binding proteins and motors, dynamically assemble and reorganize in response to a variety of biochemical and mechanical signals to produce specific architectures. This is especially true for the actin and myosin cytoskeleton, the dominant architectural element in cells [1].

The Rho family of GTPases (RhoA, Rac, and Cdc42) often act between upstream cues and downstream actomyosin re-arrangement, functioning as master regulators of the cytoskeleton [2]. The spatial and temporal activation of Rho GTPases coupled with their mutual antagonism enables critical cellular processes like migration and cell division [3, 4]. Rho GTPases signal by binding and activating downstream effectors. For example, GTP-bound RhoA can activate formins to induce actin polymerization and bind to Rho-associated coiled-coil containing protein kinase (ROCK) to phosphorylate and activate myosin 2, thereby driving actomyosin activity [5–7]. Actomyosin contractility can then actuate mechanical responses by other cytoskeletal elements and feedback to influence Rho GTPases [8, 9]. This dynamic Rho-driven mechanical crosstalk model likely influences all cytoskeletal elements, but has been largely unexplored in the septin cytoskeleton.

Septins are GTP-binding proteins that polymerize to form cytoskeletal filaments and are broadly conserved across multiple species [10–12]. In mammals there are 13 septin genes divided into four families based on sequence homology, with members of each family showing tissue specific expression [13]. Mammalian septins oligomerize into non-polar palindromic subunits consisting of either six or eight monomers drawn from three or four families, respectively [14]. These hexamers and octomers then polymerize into filaments and other higher order structures [15, 16]. First discovered as critical components of cytokinesis in budding yeast [17], septins play important roles across physiology, including cell migration [18–20], neurogenesis [21], immunology [22, 23], host-pathogen interactions [24] and ciliogenesis [25]. Furthermore, knockout of multiple individual septin genes are embryonically lethal [26–28] and changes in septin expression are linked to a number of diseases including Alzheimer’s and breast cancer [29–31], demonstrating a critical role for septins throughout cell biology.

Septins are largely considered protein scaffolds based on their ability to interact with the plasma membrane and multiple cytoskeletal proteins, in addition to their lack of associated motors. At the plasma membrane, septin distribution is heterogeneous as septins exhibit a preference for membranes with micron-scale curvature [32–34]. In vitro, septins can directly bind both microtubules and actin to induce bundling [35, 36]. In cells, however, these interactions are more complicated as septins decorate only subpopulations of both actin and microtubule networks [37]. Furthermore, a previous report suggested septins could also bind directly to non-muscle myosin 2A (hereafter, myosin 2) [38]. Together these features make septins prime conduits to translate both biochemical and biophysical signals in cells.

While pharmacological and genetic perturbations can globally perturb septin function, it is clear that septin localization in vivo is regulated both spatially and temporally. Cdc42 signaling has previously been shown to play a prominent role in septin regulation [39–43]. However, septin’s essential role in the mammalian contractile ring, where RhoA is the dominant signaling mechanism and actomyosin is the dominant architectural element, suggests septins may respond to additional biochemical inputs. Septins have also been broadly implicated in cell mechanics [44, 45], which is largely mediated by RhoA signaling, although their precise contribution remains uncertain. This link to cytokinesis and mechanics, coupled with in vitro actin binding and possible myosin 2 binding, compelled us to more carefully dissect the spatiotemporal contribution of actin, myosin 2 and RhoA in septin network formation.

Like other cytoskeletal elements, septin networks are both dynamic and heterogeneous [41, 46, 47]. Optogenetic tools, therefore, are particularly well-suited to spatiotemporally probe regulation of septin networks. These tools take advantage of light-sensitive proteins that change conformation when exposed to light of certain wavelengths [48, 49]. The improved light inducible dimerization (iLID) system is a popular version of this approach based on its specificity, rapid association and dissociation kinetics, and need for minimal intensity of light for activation [50, 51]. In particular these tools have proven adept at locally stimulating GTPase signaling by recruiting specific guanine exchange factors (GEFs) to activate RhoA, Rac, or Cdc42 [52–57].

Here we set out to investigate a role for RhoA signaling in regulating septin accumulation in interphase cells. We first demonstrate that inhibition of myosin 2, a downstream effector of RhoA, results in septin relocalization from actin stress fibers to the plasma membrane. Optogenetic recruitment of myosin 2 alone, however, is insufficient to alter septin localization. Similarly, optogenetic activation of myosin to locally increase contraction via recruitment of myosin light chain kinase (MLCK) also fails to induce septin recruitment. Local activation of RhoA, however, results in a local increase of septin in an anillin independent manner. Combined, these data suggest that RhoA signaling, and not just contractility, is able to induce septin assembly, expanding the reach of septins in these critical biochemical signaling networks.

## Results

### Local recruitment of myosin 2 is not sufficient to recruit septins

As septins and myosin 2 colocalize at the contractile ring and have also been previously reported to interact [38] we set out to test whether inhibiting myosin 2 would impact septin localization. We found that treating fibroblasts with the myosin 2 inhibitor blebbistatin resulted in disassembly of central septin networks and increased accumulation of septins at the plasma membrane (Fig. 1). As this implies that myosin may regulate septin localization, we set out to directly test whether recruitment of myosin 2 would result in increased septin accumulation. We recently developed an iLID-based tool to use light to locally recruit myosin 2 [58]. Briefly, this iLID optogenetic system consists of a transmembrane anchor (Stargazin) attached to a light-oxygen-voltage sensing domain (LOV2) whose J*α* helix has been modified with a small bacterial peptide (SsrA) which remains masked in the dark state [50, 51] (Fig. 2A). Upon blue light (*λ <* 500*nm*) exposure, the J*α* helix unfolds, exposing the SsrA peptide and allowing it to bind it’s complementary binding partner SspB attached to non-muscle myosin 2A (NM2A; Fig. 2A) [58]. Consequently, local illumination in a region spanning the cell resulted in a significant increase in myosin intensity that abated when the stimulating light was removed (Fig. 2B-E, Video 1). However, we saw no appreciable change in septin intensity or localization in response to the increasing density of myosin (Fig. 2B-E, Video 1).

**Fig. 1.**
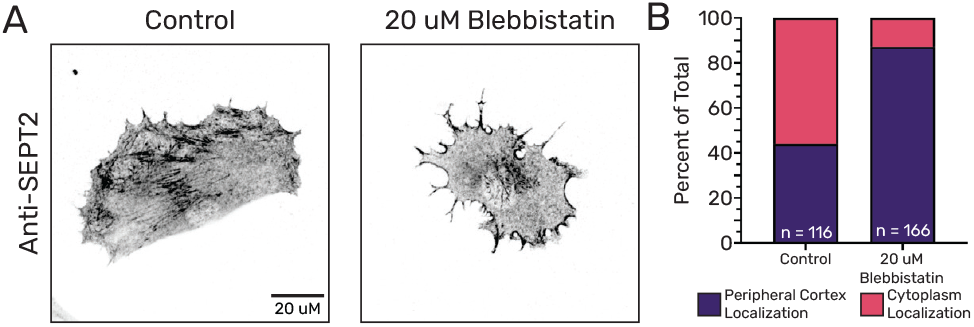
Inhibition of myosin phosphorylation results in increased septin localization on membrane. **(A)** Representative fibroblast cells treated with vehicle control or 20 *µ*M (-)-Blebbistatin for 1 hr prior to being fixed and stained with antibody against SEPT2. **(B)** Bar graph showing percent of cells with the brightest 2% pixels overlapping with the peripheral cortex. (purple) with control (n = 116) and 20 *µ*M (-)-Blebbistatin (n = 166) treatment.

**Fig. 2.**
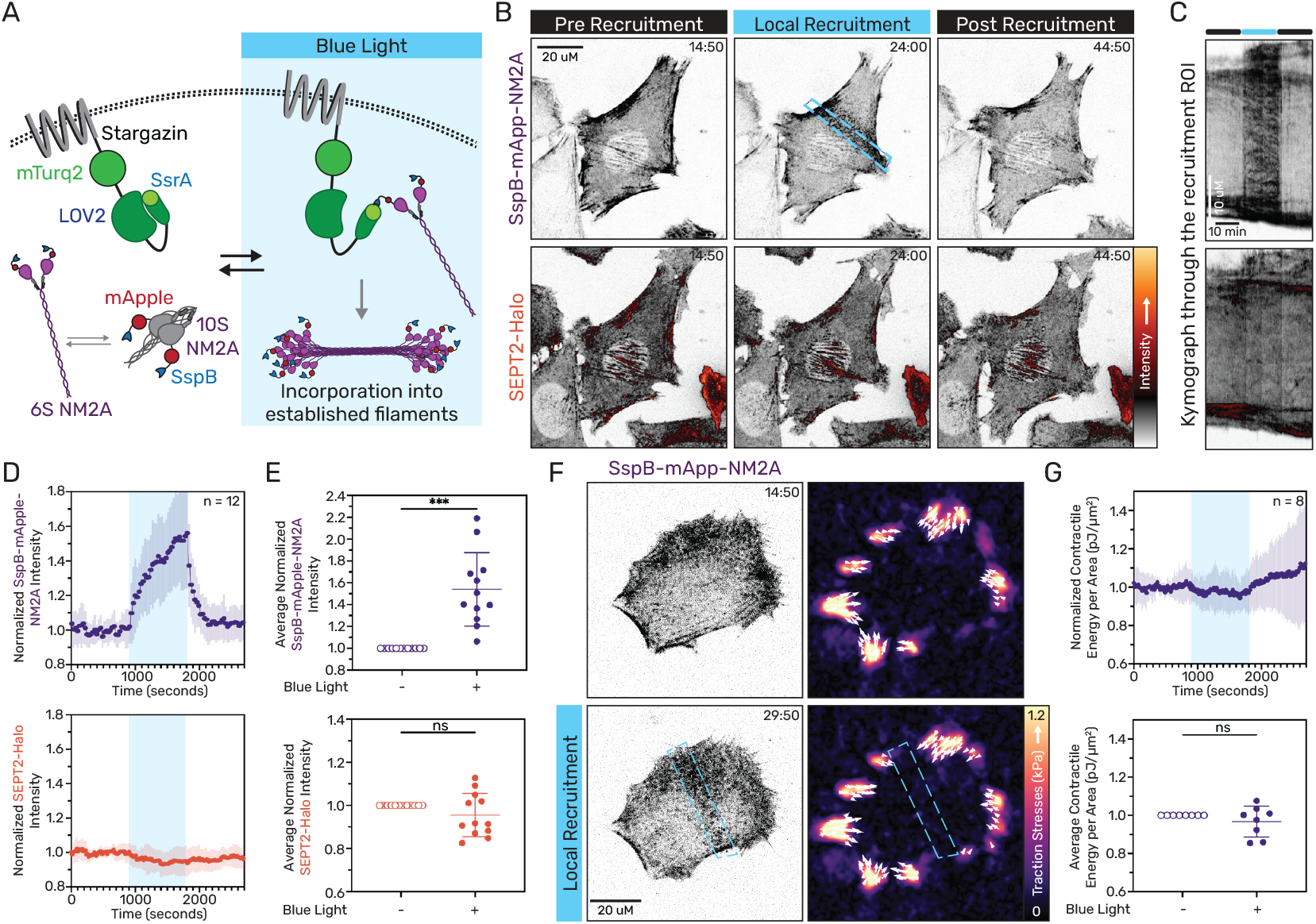
Local recruitment of myosin 2 is not sufficient to recruit septins. **(A)** A schematic of the tool to locally recruit myosin. A LOV2-SsrA molecule and mTurq2 fluorophore are anchored to the plasma membrane via stargazin. Simultaneously, an NM2A construct tagged with an mApple-SspB is expressed in the cytosol. Upon local blue light stimulation (blue box) the SspB-mApple-NM2A is recruited to the plasma membrane where it can enhance filament assembly. When blue light is removed, the two components disassociate. **(B)** Representative images of a Halo-labeled SEPT2-Halo-KI fibroblast expressing the SspB-mApple-NM2A, immediately prior to, during, and after local recruitment (blue dashed box; see Video 1). **(C)** A kymograph drawn through the activation region shows increased NM2A intensity but no change in SEPT2 intensity during the activation period. **(D)** Average SspB-mApple-NM2A and SEPT2-Halo intensities (mean *±* SD) over time in the recruitment region, normalized to the pre-recruitment time points. Recruitment period is indicated by the blue background. **(E)** Scatter plots of average intensity changes in the recruitment region between the 50 s pre-recruitment and the final 50 s of recruitment per cell. **(F)** Representative images of myosin and the accompanying traction stress maps prior to and during local myosin recruitment (see Video 2). **(G)** Average measurements of the normalized total contractile work per area performed by cells over time, and scatter plots comparing the contractile work per area between the 50 s pre-activation and the final 50 s of activation per cell. * and *** indicate p ≤ 0.05 and ≤ 0.001, respectively. n values indicate the number of cells analyzed.

Interestingly, while we previously found myosin 2 recruitment self-stimulated myosin filament assembly in the lamella where myosin filaments were initially sparse [58], here we saw an increase in intensity of myosin but minimal changes in architecture when recruiting to regions where myosin 2 filaments were already abundant. To check whether the increase in myosin impacted the overall contractility of the cell, we used traction force microscopy (TFM) to measure cellular force production. As forces generated in the cytoskeleton are transmitted to the substrate via focal adhesions, local changes in contractility in the cytoskeleton can alter traction stress measurements outside the region of recruitment [59]. We therefore measured the total contractile work performed by the cell normalized by the cell area, which we have shown previously is a better reporter of cell contractility [60]. Interestingly, we found that despite the increase in myosin in these regions, the total contractile work performed did not change (Fig. 2F-G, Video 2). Together, these data indicate that myosin alone is insufficient to induce septin accumulation.

### Locally inducing contraction does not alter septin localization

While our data suggest that myosin is not directly interacting with septins, the septin rearrangement we observed upon myosin inhibition (Fig. 1) suggests that myosin contractile activity could still aid in septin localization. We therefore sought to increase local myosin contractile activity by modulating its phosphorylation. To achieve this we swapped the stargazin anchor in our iLID system with myosin regulatory light chain (RLC), and attached the SspB to the kinase domain of myosin light chain kinase (MLCK) (Figs. 3A, S1, Video 3). This tool should therefore stimulate both RLC Thr18/Ser19 phosphorylation on myosin monomers to induce nascent filament formation, and simultaneously increase the RLC phosphorylation in existing filaments to preserve/prolong their activity (Fig. 3A). To validate this approach, we again measured the total contractile work performed by the cell using TFM (Fig. 3B, Video 4). We saw a steady increase in traction forces and the overall contractile work performed by the cell in response to local recruitment of MLCK to myosin, which decreased as the stimulating light was removed (Fig. 3C). This demonstrates that recruitment of the kinase domain of MLCK to myosin results in increased contractility.

**Fig. 3.**
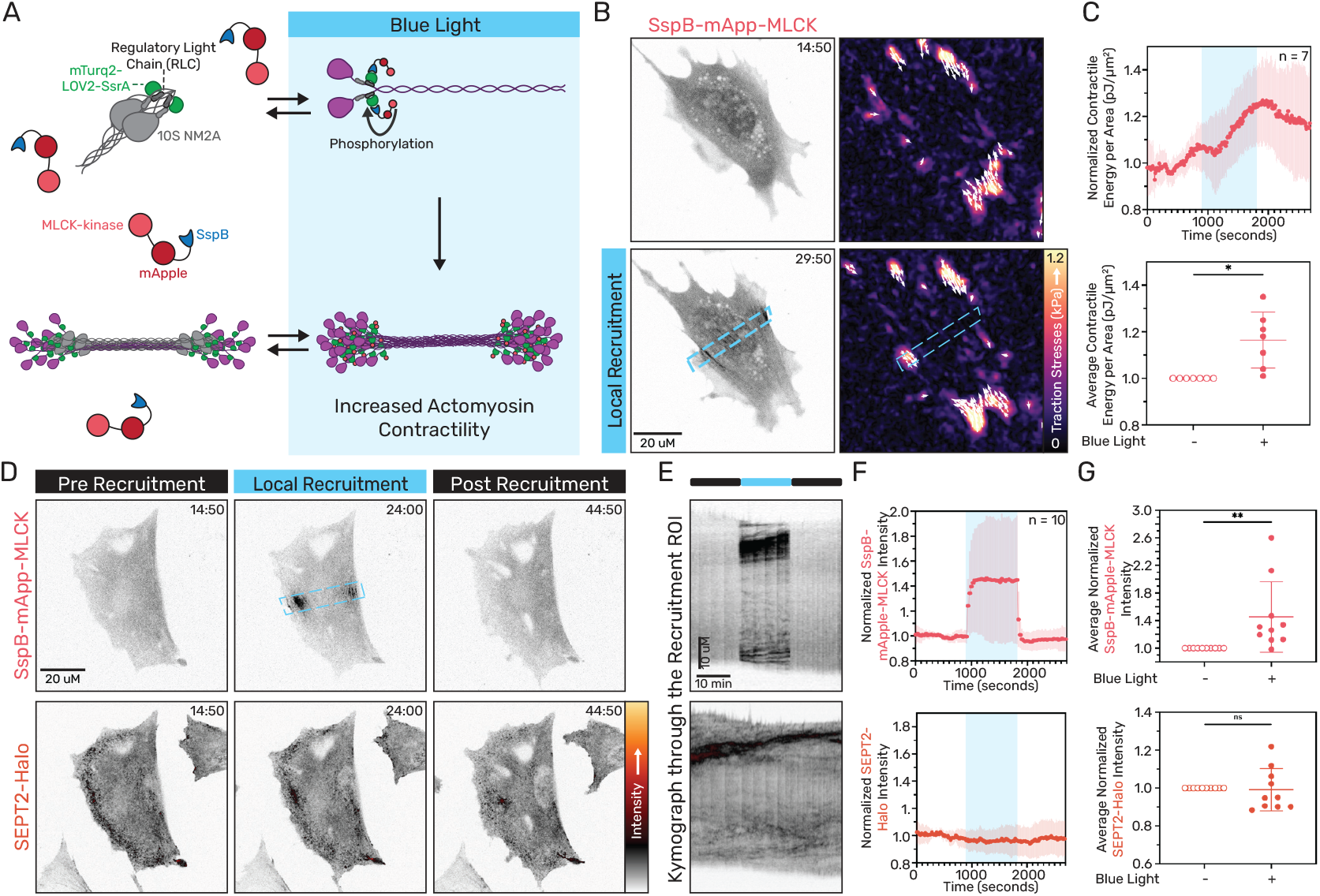
Local recruitment of MLCK to myosin filaments elevates contractility but does not alter septin localization. **(A)** A schematic of the optogenetic MLCK tool used to increase myosin-based contractility. The LOV2-SsrA molecule was anchored to the myosin RLC while the kinase domain of MLCK tagged with SspB-mApple was expressed in the cytosol. Upon blue light illumination, SsrA-SspB binding should result in increased activation of myosin by the recruitable MLCK. When the blue light is turned off, MLCK dissociates from the myosin RLC. **(B)** Representative images of a fibroblast expressing the Sspb-mApple-MLCK and the corresponding traction maps prior to and during local recruitment (blue dashed box; see Video 4). **(C)** Average measurements of the normalized total contractile work per area performed by cells, and scatter plots comparing the contractile work per area before and during activation (mean ± SD). **(D)** Representative images of a SEPT2-Halo-KI fibroblast expressing SspB-mApple-MLCK immediately prior to, during, and after local stimulation (blue dashed box; see Video 5). **(E)** Kymograph drawn through the region of recruitment in the cell shown in (D). **(F)** Normalized SspB-mApple-MLCK (top) and SEPT2-Halo (bottom) intensities (mean ± SD) over time in the region of recruitment, with recruitment periods indicated by the blue background. **(G)** Scatter plots of average intensity changes in the recruitment regions between the 50 s pre-recruitment and the final 50 s of recruitment per cell (mean ± SD). ns, *, and ** indicate p *>* 0.05, ≤ 0.05, and ≤ 0.01, respectively. n values indicate the number of cells analyzed.

While the traction stresses largely increased outside the region of MLCK recruitment due to tension propagating along actin fibers to focal adhesions, inside the recruitment region we saw a local increase in signal of the Sspb-MLCK (Fig. 3B-G, Video 5). This indicates that within the cytoskeleton, contractility and tension should be increasing locally within the recruitment region where the MLCK is acting on myosin. To test whether an increase in local contraction impacts septins, we repeated our local MLCK recruitment while visualizing septins. We found that increasing contractility was not sufficient to drive septin recruitment, as we did not see any significant changes in septin intensity in the region of MLCK recruitment (Fig. 3D-G, Video 5). Together these data suggest that myosin-based contractility is not sufficient to recruit or alter septin localization in interphase cells.

### Local recruitment of LARG-DH specifically activates RhoA

Since increased myosin accumulation and contractility alone are insufficient to recruit septins, additional signaling must be necessary. Septins are prominently found in the contractile ring during cytokinesis [61]. In this critical structure there is both local elevation of RhoA signaling [52], and an increase in anillin, a scaffolding protein which can bind both septin and active RhoA [62]. We thus set out to investigate whether these proteins contribute to septin organization and accumulation in interphase cells.

RhoA is regulated by the activity of multiple guanine-nucleotide exchange factors (GEFs) and GTPase activating proteins (GAPs), which can target either individual or multiple RhoGTPases [63]. Multiple optogenetic probes to control RhoA activation have been developed, targeting either specific GEFs or RhoA itself. Leukemia associated Rho GEF (LARG; also known as ARHGEF12) is a RhoA specific GEF [64]. Previous works have shown that optogenetically recruiting the DH domain from LARG is sufficient to activate RhoA locally and stimulate downstream actin polymerization, myosin phosphorylation, and force production (Fig. S2A-B) [52, 56, 59, 65]. As previous studies have shown that Cdc42, another Rho GTPase, and its downstream effectors can regulate septin formation [39, 66], we first confirmed that the truncated LARG-DH maintained its specificity for RhoA and did not also activate Cdc42. Using an iLID system again anchored to the plasma membrane, we locally recruited SspB-LARG-DH while simultaneously using biosensors to measure either RhoA (Fig. S2C-F, Video 6) or Cdc42 activity (Fig. S2G-J, Video 7). While recruitment of LARG-DH reliably increased local RhoA activity, we saw no similar increase in Cdc42 activity (Fig. S2C-J), indicating that the truncated LARG retains its specificity for RhoA.

### Local RhoA activation does not result in anillin recruitment

We next investigated whether local RhoA activation results in recruitment of anillin, which plays a key role in coordinating the actomyosin contractile ring formation [62]. Anillin contains a Rho binding domain, a septin binding domain, a pleckstrin homology (PH) domain on its C-terminus, and myosin and actin binding domains on its N-terminus [62]. While anillin is thought to reside primarily in the nucleus during interphase, recent results have suggested that it may have cytoplasmic roles outside of cell division [67]. We first fixed and stained cells to compare anillin expression between cells in interphase and during cytokinesis. The residual anillin expression in interphase relative to cytokinesis was primarily restricted to the nucleus (Fig. 4A). To test whether local recruitment of RhoA resulted in increased anillin recruitment in interphase cells, we locally activated RhoA and fixed after 15 minutes of activation, a time point consistent with our other optogenetic experiments (Fig. 4B). Anillin signal remained unchanged in interphase cells where local RhoA was activated (Fig. 4B), from which we conclude that interphase recruitment of LARG-DH activates RhoA without significantly modulating any cytoplasmic anillin.

**Fig. 4.**
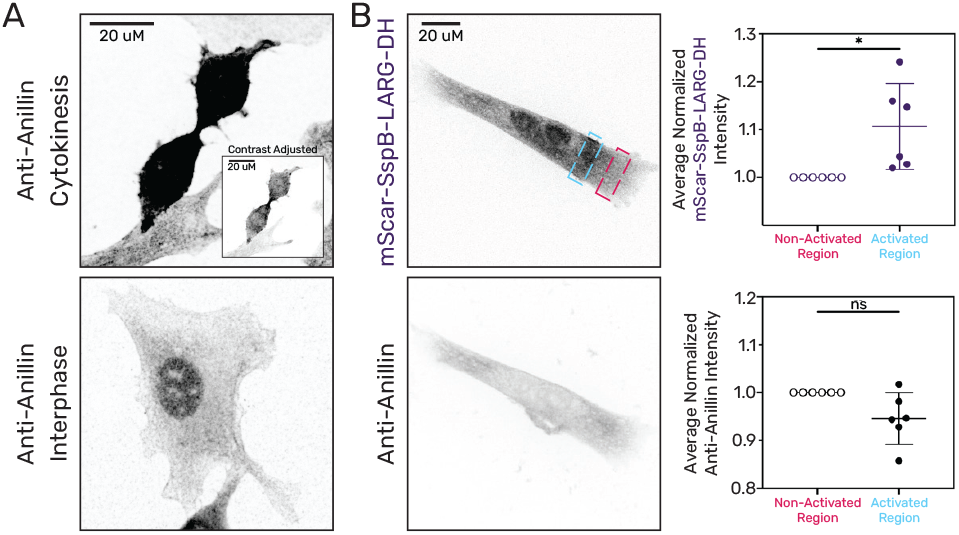
Local RhoA activation does not result in anillin recruitment. **(A)** Fibroblasts were fixed and stained with an anti-anillin antibody during cytokinesis and interphase. Images are shown with identical contrast settings. Inset is a contrast adjusted image to show that that anillin expression is increased at the midbody as expected during cytokinesis. **(B)** Representative fibroblast stably expressing Stargazin-mTurq2-LOV2-SsrA and transiently expressing mScarlet-SspB-LARG-DH (top), fixed during local activation of RhoA and stained with anti-Anillin (bottom). **(C)** Scatter plots showing differences in average fluorescence intensity between the light-activated region (blue box) and an adjacent non-activated region (red box). ns and * indicate p *>* 0.05 and ≤ 0.05, respectively.

### Local RhoA activation induces septin recruitment

Having confirmed that local recruitment of LARG-DH should activate only RhoA, we set out to investigate whether RhoA activation would impact septin localization. To test this, we locally activated RhoA and measured changes in SEPT2 intensity in the region of activation (Fig. 5A-D). In contrast to the mScar-LARG-DH which shows an immediate increase, we routinely saw that activation of RhoA resulted in a slower accumulation of septin into the activation region (Fig. 5A-C, Video 8). During the 15 min activation period septin intensity progressively and significantly increased by ~20% and decreased when the stimulating light was turned off (Fig. 5C-D). Since RhoA activation leads to downstream actin polymerization and myosin phosphorylation, which combine to increase force production, we sought to test whether activation of RhoA resulted in a significantly larger increase in contractility than the local recruitment of MLCK. Using TFM, we locally activated RhoA and observed an ~15-20% increase in the contractile work being performed by the cell (Fig. 5E-F, Video 9), similar to the increase we observed when locally recruiting MLCK (Fig. 3C). Together, these data suggest that local RhoA activation recruits septins through a mechanism specific to RhoA and its downstream effectors, rather than simply by increasing contractility.

**Fig. 5.**
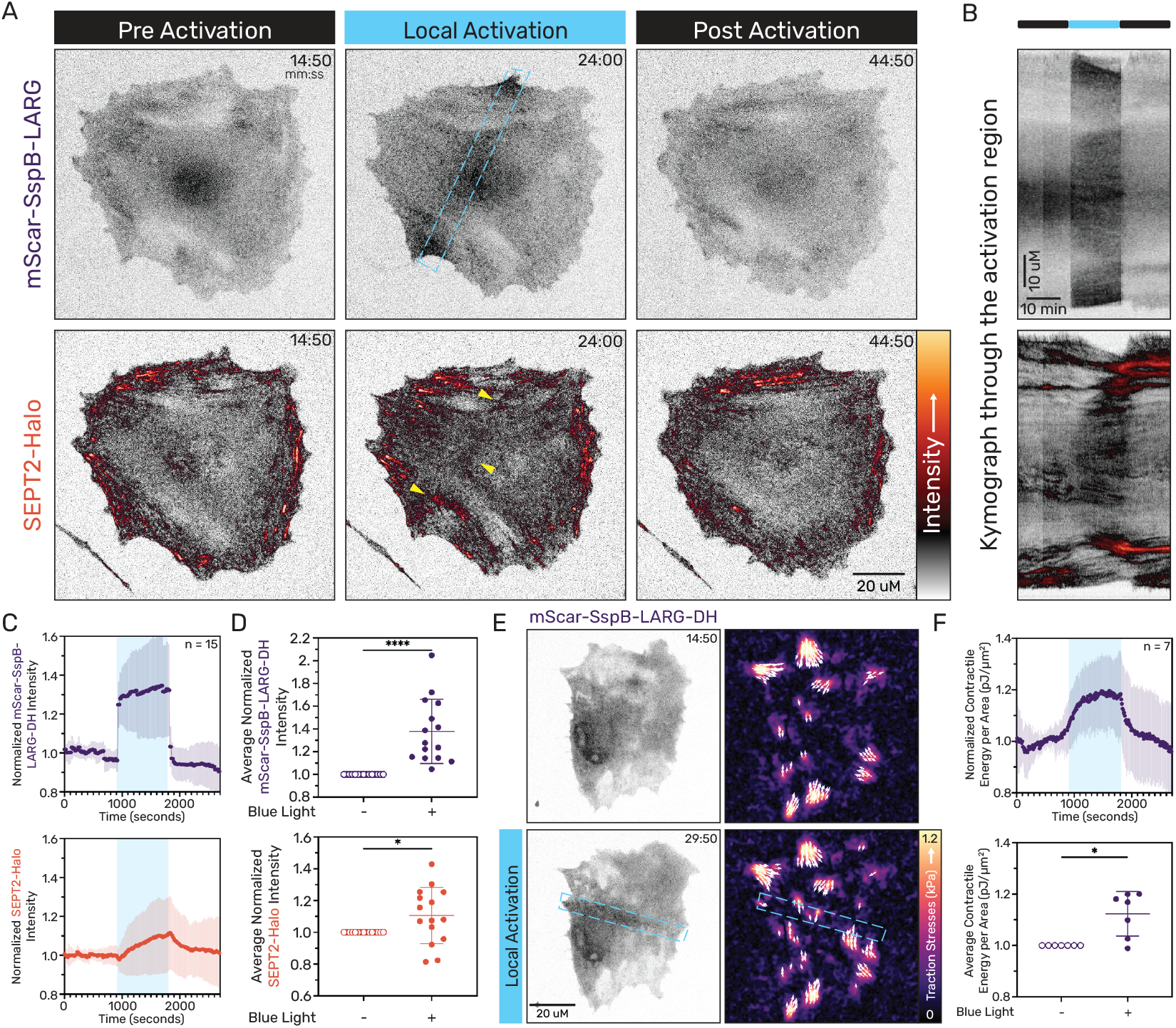
Local RhoA activation induces septin accumulation. **(A)** Representative images of SEPT2-Halo-KI fibroblast, stably expressing Stargazin-mTurq2-LOV2-SsrA and transiently expressing mScarlet-SspB-LARG-DH. Local blue light stimulation (blue dashed box, top row) induces rapid mScarlet-SspB-LARG membrane recruitment, followed by septin accumulation. Both signals dissipate when blue light is turned off (see Video 8). **(B)** A kymograph through the activation region shows increased LARG and SEPT2 intensities during the activation period. **(C)** Normalized mScarlet-SspB-LARG and SEPT2-Halo intensities (mean ± SD) in the region of RhoA activation are plotted over time, with activation periods indicated by the blue background. **(D)** Scatter plots of average intensity changes in the activation region between the 50 s pre-activation and the final 50 s of activation per cell. **(E)** Representative images of a fibroblast expressing the mScar-SspB-LARG-DH and the associated traction maps prior to and during local activation of RhoA (blue dashed box; see Video 9). **(F)** Average measurements of the normalized total contractile work per area performed by cells, and scatter plots comparing the contractile work per area in the before and during activation (mean ± SD). * and **** indicate p ≤ 0.05 and ≤ 0.0001, respectively. n values indicate the number of cells analyzed.

## Discussion

While the role of septins in cell division has been clearly defined, their functions during the rest of the cell cycle are more varied. It has long been reported that septins colocalize and interact with the actomyosin cytoskeleton [15], leading to suggestions that septins are responding to myosin generated forces [44]. Our data showed that inhibition of myosin causes septins to relocalize from stress fibers to the cortex (Fig. 1), but that reciprocal experiments increasing myosin concentration (Fig. 2) or contractility (Fig. 3) are insufficient to alter septin filament localization. In contrast, local activation of RhoA robustly promoted septin recruitment at the site of activation (Fig. 5) in an anillin independent manner (Fig. 4). These data highlight that RhoA activity provides an alternative signaling pathway to lead to septin recruitment.

Given septin’s appearance on stress fibers and other actin structures in the cell [15, 19, 47, 68], the potential for crosstalk between septin and myosin has been a tantalizing hypothesis. Previous work suggested that SEPT9 and myosin may compete for similar actin binding sites [69], while other work suggested that SEPT2 can directly bind myosin [38]. While it is clear that septin and myosin can be in close proximity on actin filaments, our data do not support either a direct interaction or competition for sites, as we see neither an increase or decrease in septin intensity. As these previous studies both used purified proteins, it is possible that in the cell the presence of additional actin binding proteins and signaling partners limits their direct interactions. This would be consistent with the presence of anillin to act as scaffold for these two proteins during cell division [62], and with other recent work showing that Bni5 in yeast performs a similar function [70]. It is also possible that septin and myosin are regulated independently despite their colocalization, as was recently demonstrated in Drosophila border cell clusters [20].

On the signaling side, Rho family GTPases are obvious candidates to contribute to septin regulation given their strong impact on actin dynamics and architectures. Previous works established that Cdc42 and its downstream effectors, BORGs, can impact septin behavior [39, 40, 42, 66, 71]. Given the prominent role of both RhoA and septins in cell division, however, the possibility of a RhoA-septin signaling axis seems reasonable. While our findings do not determine if septins and RhoA directly interact or require intermediate effectors, we found consistent septin accumulation in regions of local RhoA activation. These data fit with other recent findings of RhoA regulating septin accumulation at the membrane of migrating border cells in Drosophila [20], and with SEPT9 activating RhoA at sites of mitochondrial fission [72]. Together, these findings suggest a more extensive role for RhoA in regulating septins.

The specific mechanism by which RhoA influences septin organization remains unclear. Our MLCK data indicates that downstream force production alone is insufficient (Fig. 3). While RhoA stimulates actin polymerization via formins (Fig. S2B), septins only decorate a subpopulation of stress fibers in the cell (Fig. 1) and are often absent from other regions with increased actin polymerization, such as the lamellipodia or filopodia. Therefore, actin polymerization alone also appears insufficient for septin recruitment and additional signaling components are likely required. The fact that the septin response builds slowly during local activation hints that it is likely not a direct interaction with RhoA, as the recruitment of the GEF happens almost immediately (Fig. 5C). It is possible that the combination of RhoA-driven actin polymerization and myosin 2 contractility create a contractile network that is favorable for septin enrichment, consistent with the parallels drawn to the contractile ring. More probably, additional downstream RhoA effectors are aiding in septin recruitment, as is the case with Cdc42. Septins have been identified in a number of proteomic screens related to focal adhesions and other actomyosin structures [73–76], and future work will be required to whittle these lists down to a manageable number of potential targets.

In summary, this work demonstrates the power of optogenetic switches and activation in deciphering biochemical signaling within cells. Our data reveals a role for RhoA in reorganizing septin localization in response to RhoA activity outside of cell division. Finally, while septins do not appear to be directly responding to increases in myosin activity or contractility, their positioning on actin stress fibers and dynamic ability to reorganize in the cytoskeleton suggests that they may have other potential roles to play in mechanotransduction.

## Materials and Methods

### Cell culture and transfection

Mouse embryonic fibroblasts (MEFs) were a kind gift of Mary Berckerle’s laboratory (University of Utah, Salt Lake City, UT). HEK293T cells (CRL-3216; ATCC) were used to prepare lentivirus. Cells were cultured in Dulbecco’s modified Eagle’s medium (10013CV; Corning), supplemented 10% FBS (10437-028, Gibco), 1% antibiotic-antimycotic (30004CI; Corning) and 5 µg/mL prophylactic plasmocin (ant-mpp; Invivogen) at 37°C in 5% CO^2^. All cells were grown on uncoated plastic tissue culture dishes and plated on #1.5 glass bottom dishes (Cellvis) or coverslips (Corning)for imaging.

At 16-20 hrs before LARG-DH and 40-44 hrs before NM2A and MLCK experiments, 100-125k MEF cells were transfected with the relevant plasmids, using the Neon Electroporation system (Thermo Fisher Scientific) with 3 × 20 ms pulses of 1000V and 4 µg total DNA in a 100 µL reaction. Cells were incubated with 100 nM Janelia Fluor JFX554 (HT030; Promega) or Janelia Fluor 646 (HT106A; Promega)/JFX650 HaloTag ligand (HT1070; Promega) overnight and washed out three times with PBS before imaging in media to visualize RhoA biosensor and Halo-tagged septin, respectively.

### Generation of CRISPR knock-in cell lines

Halo-SEPT2 knock-in cells were derived from parental fibroblasts using CRISPR/Cas9. We generated pSpCas9(BB)-2A-Puro (PX459) V2.0 (Plasmid #62988; Addgene) with target sequence 5’-AAACTTCATCAATAACCCGC-3’using established protocols [77]. To generate donor plasmids, pUC57 was digested with EcoR1 and Stu1 and purified. A four-piece Gibson assembly was then performed with three gBlocks (IDT): (1) a 794 bp 5’HDR of genomic sequence immediate upstream of the endogenous start codon, (2) HaloTag with an 18 amino acid GS-rich linker, (3) an 802 bp 3’HDR of genomic sequence immediately downstream of the endogenous start codon with silent PAM site mutation. Fibroblasts were transfected with donor and target-Cas9 plasmids and single-cell sorted. Individual clones were evaluated for knock-in via Western blotting and microscopy.

### Expression vectors and molecular cloning

pSspB-mApple-NM2A and pSspB-EGFP-LARG-DH were previously described [58, 78]. pdimericTomato-1xwGBD was a gift from Dorus Gadella (Plasmid #176099; Addgene) [79]. pShuttle-CMV-Halo-AHPH (“RhoA biosensor”) was generated by digesting pShuttle-CMV (Plasmid #16403; Addgene) [80] with Not1 and inserting PCR-amplified Halo-Tag and AHPH from pEGFP-RhoA Biosensor (Plasmid #68026; Addgene) [81] using Gibson cloning. pmScarlet-SspB-LARG-DH was generated in two steps using pLV-GFP (Plasmid #25999; Addgene) [82] as a backbone. We used Gibson cloning to sequentially replace GFP with mScarlet, then inserted SspB-LARG-DH amplified from mCherry-NES-SspB-LARG-DH-P2A-iLIDcaax (Plasmid #173870; Addgene) [56]. To generate pSspB-LARG-iRFP720 plasmid, SspB-LARG was inserted into piRFP720-N1 vector (Plasmid #45461; Addgene) [83] using the AgeI and HindIII restriction sites. To generate a photorecruitable MLCK-kinase (pSspB-mApple-MLCK-kinase), we used Gibson cloning to insert human MLCK (NM 053025.4) catalytic domain corresponding to AA G1424 - K1770 into pSspb-mApple-NM2A [58] following removal of NM2A with HindIII and Spe1. The linker between mApple and MLCK catalytic domain is YSDLELKL. To generate a myosin RLC iLID anchor (pLV-MYL12B-mTurqoise2-iLID) we used Gibson cloning to replace ADRB2 in pLV-ADRB2-mTurqoise2-iLID (Plasmid #161002; Addgene) using BamH1 restriction sites with human MYL12B coding sequence from NM 033546.4. All constructs were verified by full-plasmid sequencing.

### Stable cell lines

HEK293T cells were transfected using LipoD293 (SL100668; Signagen) and the accompanying lentivirus generation transfection protocol. Briefly, HEK293T cells were plated in 6-cm dishes and grown to 80–90% confluence. Approximately 1 hr before transfection, media was changed. Transfection complex was created with LipoD293, packaging plasmid psPax (Plasmid #12260; Addgene), envelope plasmid pmD2.G (Plasmid #12259; Addgene), and lentiviral construct pLV-Stargazin-mTurquoise2-iLID (Plasmid #161001; Addgene), and added drop-wise to the dish. Media was changed 24 hrs after transfection and collected at 48 and 72 hrs. Viral media was centrifuged at 1,600 g for 5 min. A 50% confluent 6 cm of cells (wild-type and septin knock-in MEFs) were infected with viral media supernatant. Viral media was removed after 48 hrs. At 3 days post viral infection, positive cells were selected using 2 µg/mL blasticidin (ant-bl-1; Invivogen).

### Live-cell imaging

Cells were rinsed with PBS before imaging and replenished with fresh culture media. All imaging was performed at 37°C with 5% CO2 on an Axio Observer 7 inverted microscope (Zeiss) attached to a W1 Confocal Spinning Disk (Yokogawa) with Mesa field flattening (Intelligent Imaging Innovations), a motorized X,Y stage (ASI), and a Prime 95B sCMOS (Photometrics) camera. Illumination was provided by a TTL-triggered multifiber laser launch (Intelligent Imaging Innovations) consisting of six diode laser lines (405, 445, 488, 514, 561, and 640 nm) and all matching requisite filters using a 63X, 1.4 NA Plan Apochromat objective (Zeiss). Temperature and humidity were maintained using a Bold Line full enclosure incubator (Oko Labs). The microscope was controlled using Slidebook 6 Software (Intelligent Imaging Innovations). All imaging was performed as single confocal slices. Images were captured every 10 s, except for SspB-mApple-MLCK, mScarlet-SspB-LARG-DH, iRFP720-SspB-LARG-DH, and Sspb-mApple-NM2A constructs, which were imaged every 30 s to limit photo-bleaching. Images displayed as inverted grayscale or with custom look up table (“MQ div-autumn”) [58] downloadable at https://sites.imagej.net/NeuroCyto-LUTs/ and modifiable via https://github.com/m-a-q.

### Fixed imaging

#### Blebbistatin treatment

MEFs were plated on #1.5 glass coverslips (Corning) 24 h prior to treatment. Cells were treated with vehicle or 20 *µ*m (-)-blebbistatin (856925-71-8; Cayman Chemical) for one hour. After treatment, the cell culture media was gently rinsed with cytoskeleton buffer (CB) (19.52 g MES [0.1 M], 6.10 g MgCl2 [0.03 M], 102.88 g KCl [1.38 M], 7.6 g EGTA [0.02 M])and replaced with a fixing and blocking solution (0.15 g BSA, 2.5 ml 16% paraformaldehyde solution, 50 *µ*l Triton, and 7.5 ml CB) for 1 hour at room temperature on a rocker. Cells were washed 3×5 min with 1x PBS before incubating overnight at 4° in primary antibody for SEPT2 (11397-1-AP; Proteintech) diluted 1:300 in antibody solution (0.15 g BSA, 50 *µ*l Triton, and 10 ml CB). The next day, coverslips were washed 3×5 min with 1x PBS before incubating for 1 hour in the dark in rabbit fluorescent secondary antibody (A11036; ThermoFisher Scientific) diluted 1:1000 in antibody solution with 1:1000 Atto-Phalloidin (65906-10NMOL; Sigma-Aldrich). A final PBS wash was performed (3×5 min) before mounting the cells on a glass slide with ProLong Glass Antifade Mountant (P36980; ThermoFisher Scientific). Cells were imaged at least 24 hrs after mounting. For analysis, the top 2% of SEPT2-intensity pixels were selected and membrane localization was determined in a binary manner based on whether these pixels overlapped with the cell membrane.

#### Anillin staining

Local RhoA activation was performed for 15 minutes and then ice-cold methanol was added to the coverslip within seconds after the last blue light exposure while the sample was still on the microscope stage. Fixation for 20 min at 37° was followed by three PBS washes. Permeabilization was carried out for 5 minutes in buffer containing 0.5% Triton X-100 in CB. For blocking, cells were incubated with 1.5% BSA in CB for 1 hour at room temperature on a rocker. Cells were then incubated overnight at 4°C with polyclonal anti-anillin sera (1:500; Gift of Michael Glotzer, University of Chicago). The next day, cells were washed three times with PBS and incubated with secondary antibody (goat anti-rabbit Alexa Fluor 647; 1:1000) (#A32733; ThermoFisher Scientific) for 1 hour at room temperature. All post-fixation incubations were performed on a rocker. The average intensity within the activation region was obtained for both LARG and anti-anillin and then normalized to the average intensity in an unactivated region.

### iLID photorecruitment and analysis

Before each time lapse, the potential cell was imaged with a test time lapse consisting of 640nm/561nm/445nm excitation captures taken sequentially, and then repeated after 5 s (Fig. S1). If the anchor and its binding partner were both expressed in the cell, we would see relocation of the binding partner to the anchor following the 445 nm excitation. Once expression of both components was confirmed, the cell was kept in the dark for at least 5 minutes to allow the two components to fully dissociate. During this time, a box with a width of ~5-10 *µ*m was drawn across the ventral surface of a cell in Slidebook 6. We then proceeded with steady state imaging for 5 (Figs. 4, S 2) or 15 min (Figs. 2, 3, 5), before the local recruitment region was illuminated with a 405 nm laser for the same duration (5 or 15 min) at 6 *µ*W, followed by additional period of imaging (5 or 15 min) without local stimulation. Illumination of the local recruitment region was repeated every 10 s prior to the imaging at that time point. Cell counts are provided in the figures.

Analysis was performed using Python. Cell masks for each cell were generated in FIJI by thresholding the signal in the 561 nm channel. Images were corrected for photobleaching and flat-field. Following correction, the intensity within the region of activation (region extracted from Slidebook6) and cell mask was calculated for each image/frame. This intensity was normalized to the mean intensity of the frames prior to photo recruitment and plotted as normalized intensity over time. Where necessary, intensities were normalized to a non-activated region to account for shifts in focus.

### Traction force microscopy and analysis

Traction force microscopy was performed as previously described [84]. Coverslips were prepared by incubating with a 2% solution of 3-aminopropyltrimethyoxysilane (313255000; Acros Organics) diluted in isopropanol, followed by fixation in 1% glutaraldehyde (16360; Electron Microscopy Sciences) in ddH_2_O. Polyacrylamide gels (shear modulus: 8.6 kPa—final concentrations of 7.5% acrylamide (1610140; Bio-Rad) and 0.3% bis-acrylamide (1610142; Bio-Rad)) were embedded with 0.04 µm fluorescent microspheres (F8789; Invitrogen) and polymerized on the activated glass coverslips for 1 hr at room temperature. After polymerization, gels were rehydrated for at least 60 min. To cross-link the gels with fibronectin, gels were treated with the cross-linker Sulfo-Sanpah (22589; Pierce Scientific), photoactivated with UV light for 5 min and then rinsed thoroughly with ddH_2_O. Gels were then incubated with 40 µL of 1 mg/mL human plasma fibronectin (FC010; Millipore) for 1 hr at room temperature. Following fibronectin cross-linking, transfected cells were plated on the gels and allowed to spread overnight. Images were taken of both the cells and the underlying fluorescent beads. Following imaging, cells were detached from the gel using 0.05% SDS (L22010; Research Products International Corp) and a reference image of the fluorescent beads in the unstrained gel were taken.

Analysis of traction forces was performed using code written in Python (available at https://github.com/OakesLab/TFM) according to previously described approaches [84, 85]. Prior to processing, images were flat-field and photobleach corrected and aligned to the reference bead image with the cell detached. Other acquired channels were shifted using the same alignment measurements from the bead channel. Displacements in the beads were calculated using an optical flow algorithm in OpenCV (Open Source Computer Vision Library, https://github.com/itseez/opencv) with a window size of eight pixels. Traction stresses were calculated using the Fourier Transform Traction Cytometry (FTTC) approach with a regularization parameter of 7.9 * 10^−^7. The strain energy was calculated by summing one-half the product of the strain and traction vectors in the region under the cell and normalizing by the cell area as measured from the cell mask [60].

### Statistical analysis

Statistical analyses were performed on GraphPad Prism using the non-parametric Wilcoxon matched-pair signed rank test. P-values < 0.05 were considered statistically significant. Details about sample size and p-values are included in the figure legends.

## Acknowledgments

We thank the rest of the Oakes and Beach laboratories at the Loyola University Chicago, and Elias Spiliotis at the University of Virginia for many helpful discussions. This work was supported in part by NIH NIGMS grant R35-GM138183 to JRB, and NSF CAREER award #2000554 and NIH NIGMS grant R01-GM148644 to PWO.

## Author contributions

SC, JRB and PWO conceived the study. SC performed experiments and data analysis. BS, MEU, HW and JRB provided critical cell lines and reagents. SC, JRB and PWO wrote the manuscripts with critical feedback from co-authors.

