## Supplemental Figures for "Local RhoA activation induces septin recruitment"

### Supplementary Information:
### Local RhoA activation induces septin recruitment

Shreya Chandrasekar[1], Margaret E. Utgaard[1], Bradley Somerfield[1], Huini Wu[1], Jordan R. Beach[1,✉], and Patrick W. Oakes[1,✉]

[1]Dept. Cell & Molecular Physiology, Loyola University Chicago, Stritch School of Medicine, Maywood, IL 60153

### Supplemental Figures

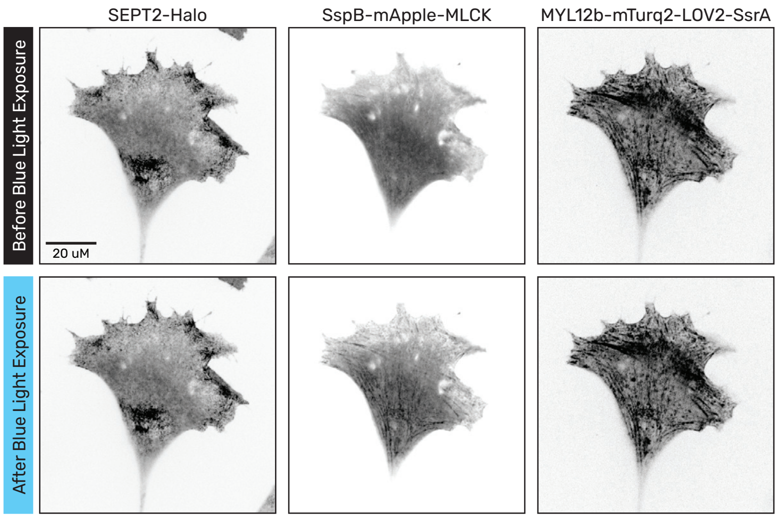


**Fig. S1. Global recruitment of MLCK to RLC.** The top row shows a SEPT2-Halo KI cell expressing the Sspb-mApple-MLCK and MYL12b-mTurq2-LOV-SsrA constructs. Images were taken in series from left to right, with a 1 s delay between the first and second rows. While no change is seen in either the SEPT2 or MYL12b proteins, the exposure to blue light causes the MLCK to bind to the MYL12b (i.e. RLC). See Video 3.

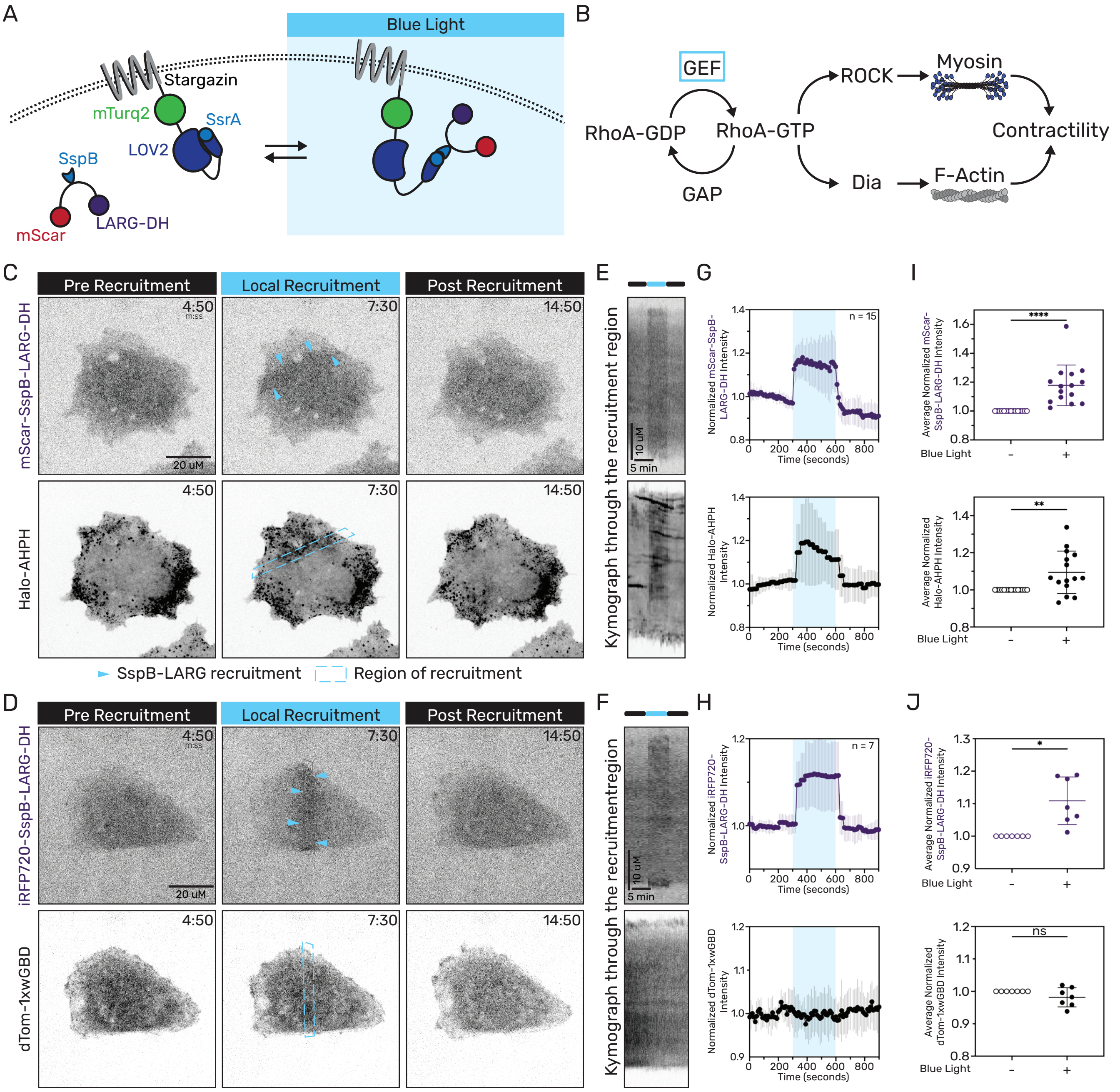


**Fig. S2. Local LARG-DH recruitment specifically activates RhoA. (A)** iLID RhoA activation optogenetic system. LOV2-SsrA is membrane-anchored via Stargazin; LARG-SspB resides in the cytosol. Blue light induces reversible SsrA–SspB binding, recruiting LARG-DH to the membrane. **(B)** RhoA pathway. GEF-mediated RhoA activation stimulates ROCK and Dia, driving myosin phosphorylation and F-actin assembly, respectively. **(C,G)** Representative images of a fibroblast stably expressing Stargazin-mTurq2-LOV-SsrA and transiently expressing photo-recruitable LARG-DH with either Halo-AHPH (C) or dTom-1xwGBD (G). Local blue light (top rows, blue box) triggers rapid SspB-LARG-DH accumulation and RhoA (C, bottom row; see Video 6) but not Cdc42 (D, bottom row; see Video 7) activation. Signals dissipate after light removal. **(D,H)** Kymographs through the activation region show increased LARG-DH and Halo-AHPH intensities (D) but no change in dTom-1xwGBD (H) during the activation period. **(E,I)** Normalized SspB-LARG-DH (top), Halo-AHPH (E, bottom), and dTom-1xwGBD (I, bottom) intensities (mean ± SD) over time in the region of activation, with activation periods indicated by the blue background. **(F,J)** Scatter plots of average intensity changes in the activation region between the 50 s pre-activation and the final 50 s of activation per cell. * $p \leq 0.05$, ** $p \leq 0.01$, **** $p \leq 0.0001$. n values indicate the number of cells analyzed.

### Supplemental Videos

**Supplemental Video 1.** Local recruitment of myosin 2 is not sufficient to recruit septins. A SEPT2-Halo-KI fibroblast expressing SspB-mApple-NM2A (left) and labeled with Halo-dye (right). Local recruitment of the SspB-mApple-NM2A is performed in the region indicated by the blue box between time points 15:00-30:00 min of the video. Time is presented as mm:ss, and the frame rate of the movie is 20 frames/sec. The movie corresponds to the data presented in Fig. 2B.

**Supplemental Video 2.** Local recruitment of myosin 2 does not alter the cell contractility. A SEPT2-Halo-KI fibroblast expressing SspB-mApple-NM2A (left) and the corresponding traction maps (right). Local recruitment of the SspB-mApple-NM2A is performed in the region indicated by the blue box between time points 15:00-30:00 min of the video. The scale of the traction maps goes from 0 to 1200 Pa. Time is presented as mm:ss, and the frame rate of the movie is 20 frames/sec. The movie corresponds to the data presented in Fig. 2F.

**Supplemental Video 3.** Global recruitment of MLCK to myoisn RLC. A SEPT2-Halo-KI fibroblast expressing SspB-mApple-MLCK is shown with and without exposure to blue light. The movie corresponds to the approach presented in Fig. S1.

**Supplemental Video 4.** Local recruitment of MLCK increases cell contractility. A SEPT2-Halo-KI fibroblast expressing SspB-mApple-MLCK (left) and the corresponding traction maps (right). Local recruitment of the SspB-mApple-MLCK is performed in the region indicated by the blue box between time points 15:00-30:00 min of the video. The scale of the traction maps goes from 0 to 1200 Pa. Time is presented as mm:ss, and the frame rate of the movie is 20 frames/sec. The movie corresponds to the data presented in Fig. 3B.

**Supplemental Video 5.** Local recruitment of MLCK is not sufficient to recruit septins. A SEPT2-Halo-KI fibroblast expressing SspB-mApple-MLCK (left) and labeled with Halo-dye (right). Local recruitment of the SspB-mApple-MLCK is performed in the region indicated by the blue box between time points 15:00-30:00 min of the video. Time is presented as mm:ss, and the frame rate of the movie is 20 frames/sec. The movie corresponds to the data presented in Fig. 3D.

**Supplemental Video 6.** Local recruitment of LARG-DH leads to local RhoA activation. A fibroblast expressing mScarlet-SspB-LARG-DH (left) and Halo-AHPH RhoA biosensor (right). Local recruitment of the mScarlet-SspB-LARG-DH is performed in the region indicated by the blue box between time points 5:00-10:00 min of the video. Time is presented as mm:ss, and the frame rate of the movie is 10 frames/sec. The movie corresponds to the data presented in Fig. S2C.

**Supplemental Video 7.** Local recruitment of LARG-DH does not alter Cdc42 activity. A fibroblast expressing iRFP720-SspB-LARG-DH (left) and dTom-1xwGBD Cdc42 biosensor (right). Local recruitment of the iRFP720-SspB-LARG-DH is performed in the region indicated by the blue box between time points 5:00-10:00 min of the video. Time is presented as mm:ss, and the frame rate of the movie is 10 frames/sec. The movie corresponds to the data presented in Fig. S2D.

**Supplemental Video 8.** Local activation of RhoA induces septin recruitment. A SEPT2-Halo-KI fibroblast expressing SspB-mApple-LARG-DH (left) and labeled with Halo-dye (right). Local recruitment of the SspB-mApple-LARG-DH is performed in the region indicated by the blue box between time points 15:00-30:00 min of the video. Time is presented as mm:ss, and the frame rate of the movie is 20 frames/sec. The movie corresponds to the data presented in Fig. 5A.

**Supplemental Video 9.** Local activation of RhoA increases cell contractility. A SEPT2-Halo-KI fibroblast expressing SspB-mApple-LARG-DH (left) and the corresponding traction maps (right). Local recruitment of the SspB-mApple-LARG-DH is performed in the region indicated by the blue box between time points 15:00-30:00 min of the video. The scale of the traction maps goes from 0 to 1200 Pa. Time is presented as mm:ss, and the frame rate of the movie is 20 frames/sec. The movie corresponds to the data presented in Fig. 5E.
